# A membrane insertion code for intrinsically disordered proteins

**DOI:** 10.64898/2026.03.12.711444

**Authors:** Fidha Nazreen Kunnath Muhammedkutty, Huan-Xiang Zhou

## Abstract

Membrane association of intrinsically disordered proteins (IDPs) mediates various cellular functions including membrane remodeling and signal transduction. Whereas membrane association through amphipathic helices and polybasic motifs is well understood, sequence determinants for the insertion of aromatic residues into the membrane hydrophobic core are still poorly characterized. Here, we decipher the sequence code for membrane insertion of aromatic-centered motifs. For an initial set of 10 9-residue aromatic-centered sequences, all-atom molecular dynamics simulations and the positioning of proteins in membranes (PPM) method produced very similar membrane insertion propensities. Applying PPM to a full library of 1.2 × 10^6^ sequences with an F, W, or Y residue flanked by L, R, G, N, or E at four positions on either side, we found that aliphatic (L) and basic (R) residues favor membrane insertion, whereas acidic (E) and polar (N) residues disfavor it. Guided by these rules, we developed a mathematical model dubbed AroMIP (<u>Aro</u>matic <u>M</u>embrane <u>I</u>nsertion <u>P</u>redictor) to predict the membrane insertion propensities of aromatic-centered motifs. AroMIP achieves 91.2%, 92.0%, and 99.7% accuracies for F-, W-, and Y-centered motifs, respectively, in disordered regions of the human proteome and is available as a web server at https://zhougroup-uic.github.io/AroMIP/. The present work provides the sequence basis and a mechanistic understanding of how IDPs employ aromatic-centered motifs to drive membrane insertion, and enriches the tools for the study of IDP-membrane association.

## Introduction

Membrane association of intrinsically disordered proteins or regions (IDPs / IDRs) mediates a host of cellular functions, including membrane remodeling,^1–6^ ion channel gating,^7–9^ signal transduction,^10–17^ membrane anchoring,^18, 19^ and host-pathogen interactions.^17, 20^ A proteome-wide screening of human transmembrane proteins found ∼60% of them to contain membrane-binding IDRs.^21^ IDRs can associate with membranes via at least three distinct modes: amphipathic helices, polybasic motifs, and aromatic-centered short motifs.^22^ Amphipathic helices most likely form upon membrane contact, resulting in a nonpolar face that projects into the acyl chain region and a polar / charged face that projects into the headgroup region or the aqueous phase. Polybasic motifs bind to the membrane surface, in particular via electrostatic interactions between R and K sidechains and phosphate groups of lipids (especially acidic lipids). Methods have been developed to predict IDP sequences that likely associate with membranes through these two modes.^7, 23^ In comparison, sequence determinants for the insertion of aromatic-centered short motifs into the membrane hydrophobic core are still poorly defined.

Aromatic residues are known to favor the membrane-water interface, based on the free energies of transfer from interface to water,^24^ the distribution of amino acids in transmembrane proteins at different positions along the membrane normal,^25–28^ and the contributions of interfacially positioned aromatic residues to the folding stability of β-barrel transmembrane proteins.^29, 30^ In the interface, aromatic sidechains can form cation-π interactions and hydrogen bonds (in the case of W and Y) with lipid headgroups and possibly hydrophobic interactions with acyl chains.^31–35^ F sidechains are also frequently positioned deeply toward the bilayer center.^28, 29^ Whereas the positions of residues in a transmembrane protein relative to the bilayer center are dictated by the three-dimensional structure of the protein, the counterparts in a peripheral membrane protein can be varied by a change in the burial depth of a local element (e.g., a loop) or an entire domain. Aromatic residues are enriched in the membrane-binding faces of peripheral membrane proteins;^36^ whether these residues insert into the acyl chain region may be context-dependent. For example, based on the effects of fluorination on membrane binding, F249 on a loop of phosphatidylinositol-specific phospholipase C penetrates into the acyl chain region, but Y258 introduced on a neighboring helix is located in the headgroup region.^37^ Cryo-electron microscopy (cryo-EM) structures of OPA1, a dynamin superfamily protein mediating fusion of the inner mitochondrial membrane, show that W775 and W778 in the flexible membrane-inserting loop (residues 768-782) are positioned deeply into the acyl chain region of cardiolipin-containing membranes.^38, 39^ Likewise, cryo-EM structures and all-atom molecular dynamics (MD) simulations suggest penetration into the acyl chain region by aromatic residues at the tip of the β1–β2 loop in gasdermins, pore-forming proteins that cause membrane permeabilization and pyroptosis (a pro-inflammatory lytic cell death).^40–42^ All-atom MD simulations also showed that W41 in a loop in the phloem lipid-associated family protein PLAT1 transiently inserts into the acyl chain region.^43^

Aromatic residues are also enriched in membrane-binding IDRs^21^ and frequently noted in the membrane binding or insertion of IDPs. For example, using magic-angle spinning ^1^H-^1^H nuclear Overhauser enhancement spectroscopy (NOESY), Zhang et al.^10^ observed strong cross peaks between acyl chains in DMPC:DMPG lipid membranes and five phenol rings from a 25-residue fragment (residues 152-176) of the IDP myristoylated alanine-rich C kinase substrate (MARCKS). Similarly, solution ^1^H-^1^H NOESY of HIV-1 trans-activator of transcription (Tat), an 86-residue IDP, in DPC micelles revealed cross peaks between acyl chains of DPC and the indole ring of the lone W residue, W11.^44^ Based on chemical shift perturbation and NMR intensity loss due to line broadening upon binding cardiolipin-containing nanodiscs, Mahajan et al.^5^ identified both an amphipathic helix and an unstructured segment (T548-A566) in the variable domain (residues 497-607) of dynamin-related protein 1 (Drp1) for membrane binding. These authors further proposed that W552 in the latter segment interacts with acyl chains, based on analogy with a WRG sequence in a transmembrane protein. A recent cryo-EM structure of Drp1 suggests membrane insertion of the variable domain.^6^ Loss of NMR intensity upon binding DMPC:DHPC or DMPG:DHPC bicelles also revealed a Y511-centered motif (and a Y483-centered motif to a lesser extent) in the pathogen-encoded signaling receptor Tir for membrane binding.^17^ It is unclear whether these Y sidechains insert into the acyl chain region. Based on paramagnetic relaxation enhancement by membrane-embedded spin labels, Jin et al.^15^ estimated that two Y residues, Y188 and Y199, in the immunoreceptor tyrosine-based activation motif (ITAM) (two Yxx(L/I) segments separated by six to eight residues) of the CD3ε component of the T-cell receptor complex had 15% to 34% probabilities of being membrane-bound. Moreover, Y199 had ∼20% probability of reaching the acyl chain region. These results stand in contrast to a static structure showing deep insertion of both Y residues solved by solution NMR spectroscopy.^45^ Lipid binding assays of fragments covering the cytoplasmic IDR of the ζ component showed that basic clusters outside the 3 ITAMs, instead of the ITAMs, bind to acidic membranes.^11^

Coarse-grained MD simulations of the T-cell receptor complex embedded in a membrane showed that Y199 of CD3ε had a propensity to penetrate into the acyl chain region; the propensity was lower for Y188 and even lower for Y residues in the 3 ITAMs of ζ chains.^46^ Also using coarse-grained simulations, Araya-Secchi et al.^14^ observed that the I_267_**F**PPV_271_ motif in the cytoplasmic IDR of the prolactin receptor interacted with acyl chains more often than with headgroups. Other aromatic-centered motifs were found to penetrate into the acyl chain region in all-atom simulations.^9, 13, 16, 20, 41, 43, 47–50^ One end of the spectrum is stable insertion, represented by the **W**_440_**FF**G**W**_444_ motif in a 57-residue IDR of Bap1, which mediates adhesion of *Vibrio cholerae* biofilms to host membranes.^20^ The other end of the spectrum is brief excursion, e.g., by Y853 in the cytoplasmic IDR of metabotropic glutamate receptor 3 (mGluR3).^16^ Somewhere in between are the **F**_448_GI_450_ motif in the disordered juxtamembrane domain (residues 440-465) of tropomyosin receptor kinase A (TrkA),^13^ V_9_**F**_10_ in the N-terminal IDR of the bacterial K^+^ channel KtrB,^9^ and W21, W25, and W29 in the N-terminal IDR of ABHD5, a regulator of lipases.^49^ Some of these simulation results have experimental support. For example, an F-to-S mutation in the **F**_448_GI_450_ motif weakened the binding of the TrkA juxtamembrane domain to acidic membrane, whereas an R-to-W mutation at position 452 strengthened membrane binding.^13^ For Bap1, a blue shift in tryptophan fluorescence upon membrane binding is consistent with a change from an aqueous environment to a hydrophobic environment for W sidechains, thereby implicating burial in the acyl chain region.^20^ Similarly, membrane binding protects W21, W25, and W29 in ABHD5 from deuterium exchange, again implicating burial.^49^ The Y853F mutation in mGluR3, introduced to deepen membrane penetration, indeed strengthened the membrane binding of residues 845-860.^16^

Determining whether a specific residue in an IDP inserts into the acyl chain region presents a major technical challenge. Experimentally, the most useful technique is NMR spectroscopy. Although both solid-state and solution ^1^H-^1^H NOESY have identified membrane-inserted residues,^10, 44^ both face difficult issues including resolution and sensitivity. All-atom MD simulations can map out the equilibrium distributions of IDP residues with respect to the membrane surface as well as the dynamic process of membrane insertion, such simulations are always open to the question of whether sampling is sufficient.^51^ The accuracy of force fields is another issue, though more so for coarse-grained simulations.

Here we address this challenge by developing a fast, sequence-based method for predicting whether aromatic-centered motifs of IDPs insert into the acyl chain region of membranes. We focus on aromatic amino acids because, as made clear by the foregoing paragraphs, they are the most frequent ones among the 20 types of amino acids found participating in deep membrane insertion. We first performed extensive all-atom MD simulations of 10 aromatic-centered peptides (9 residues each) near POPC:POPS:PIP_2_ (70:25:5 molar ratio) membranes to characterize their membrane insertion propensities. For high throughput, we used the positioning of proteins in membranes (PPM) method^52^ and obtained membrane insertion propensities very similar to those from MD simulations. We then applied PPM to an exhaustive sequence library where each of the four flanking positions on either side of an aromatic residue was filled with one of five amino acids: L, R, G, N, and E. The PPM output was well captured by a mathematical model, dubbed AroMIP (<u>Aro</u>matic <u>M</u>embrane Insertion <u>P</u>redictor), in which each flanking residue contributed a multiplicative factor to the membrane insertion score. Lastly, we trained and tested AroMIP on IDRs in the human proteome,^53^ achieving 91.2%, 92.0%, and 99.7% accuracies for F-, W-, and Y-centered motifs, respectively. AroMIP is available as a web server at https://zhougroup-uic.github.io/AroMIP/.

## Results

### All-atom MD simulations characterize membrane-bound states of aromatic-centered peptides and pathways leading to membrane insertion

We first collected 10 aromatic-centered motifs that previous studies have classified as either membrane inserters or non-inserters (Fig. 1A). This initial set includes an F-centered motif from TrkA (sequence 1), which inserted deeply into the acyl chain region in MD simulations;^13^ a W-centered motif in Drp1 (sequence 2), which was indicated by solution NMR to be an inserter;^5^ two Y-centered segments composing the ITAM in CD3ε (sequences 3 and 4), which were estimated to have 20% and 0% probabilities, respectively, of reaching the acyl chain region based on modeling paramagnetic relaxation enhancement data;^15^ and six Y-centered segments composing three ITAMs in CD3ζ (sequences 5 to 10), which did not bind to membranes according to binding assays.^11^

**Fig. 1.** Classification of aromatic-centered motifs into membrane inserters and non-inserters by MD simulations. (A) Initial dataset containing 10 sequences (sequence 1 from TrkA, 2 from Drp1, 3 and 4 from CD3ε, and 5 to 10 from CD3ζ). (B) Initial snapshot, time trace of aromatic Z_tip_, and snapshot at 980.7 ns for sequence 1. (C) Initial snapshot, time trace of aromatic Z_tip_, and snapshot at 964.3 ns for sequence 6. (B) and (C) represent simulations where the peptides are inserted or not inserted, respectively, into the membrane. Insertion means at least 80% of the frames in the last 100 ns (shaded region) have aromatic Z_tip_ at least 3.1 Å (indicated by a red dashed line) below the phosphate plane, which is set to Z = 0. Time traces are smoothed using the moving average in a 2.5-ns window. The membrane is displayed as surface, with phosphates and all atoms above in orange while those below in gray; the peptide is displayed as cartoon in cyan, with the central aromatic residue as sticks in magenta and red. (D) The number of replicate simulations, out of a total of 20, where the peptide is inserted into the membrane.

For each sequence, we generated an initial disordered conformation using TraDES^54^ and placed it near a membrane, with the central aromatic residue just above the phosphate plane. The peptide-proximal leaflet was composed of 70% POPC (zwitterionic), 25% POPS (net charge −1), and 5% PIP_2_ (net charge −4). MD simulations were conducted in 20 replicates, each for 1 µs. Figure 1B presents a simulation where the central aromatic sidechain (sequence 1), after ∼90 ns, is stably inserted into the acyl chain region. We use a value of −3.1 Å (indicated by a red dashed line), corresponding to the mean position of glycerol C2 carbon atoms, for the Z coordinate of the aromatic tip carbon atom (Z_tip_) to demarcate insertion into the acyl chain region. In contrast, Fig. 1C presents a simulation where the peptide (sequence 4) moves away from the membrane.

We label a simulation as membrane-inserting if >80% of the frames in the last 100 ns have Z_tip_ ≤ −3.1 Å. Using this cutoff, the F- and W-centered peptides (sequences 1 and 2) have 3 and 8 simulations, respectively, out of a total of 20 each, as membrane-inserting (Fig. 1D). Among all their respective 20 simulations, the F- and W-centered peptides have 20.6% and 56.6%, respectively, membrane-inserting frames (Table 1). By contrast, none of the 20 simulations of each of the six Y-centered peptides (sequences 3-10) is membrane-inserting. We classify sequences 1 and 2 as membrane inserters (with at least one membrane-inserting simulation) whereas sequences 3-10 as non-inserters. Sequence 3 is special among the non-inserters: in two of its simulations, the percentage of membrane-inserting frames is substantial, at 57.0% and 22.4%, though not reaching the 80% threshold (to be called “partly inserting”). Correspondingly, the overall percentage of membrane-inserting frames in 20 simulations of this peptide is the highest, at 4.5%, among all six Y-centered peptides (Table 1). These outcomes are in complete agreement with the previous characterizations of the membrane-insertion propensities of these peptides.^5, 11, 13, 15^

**Table 1.**
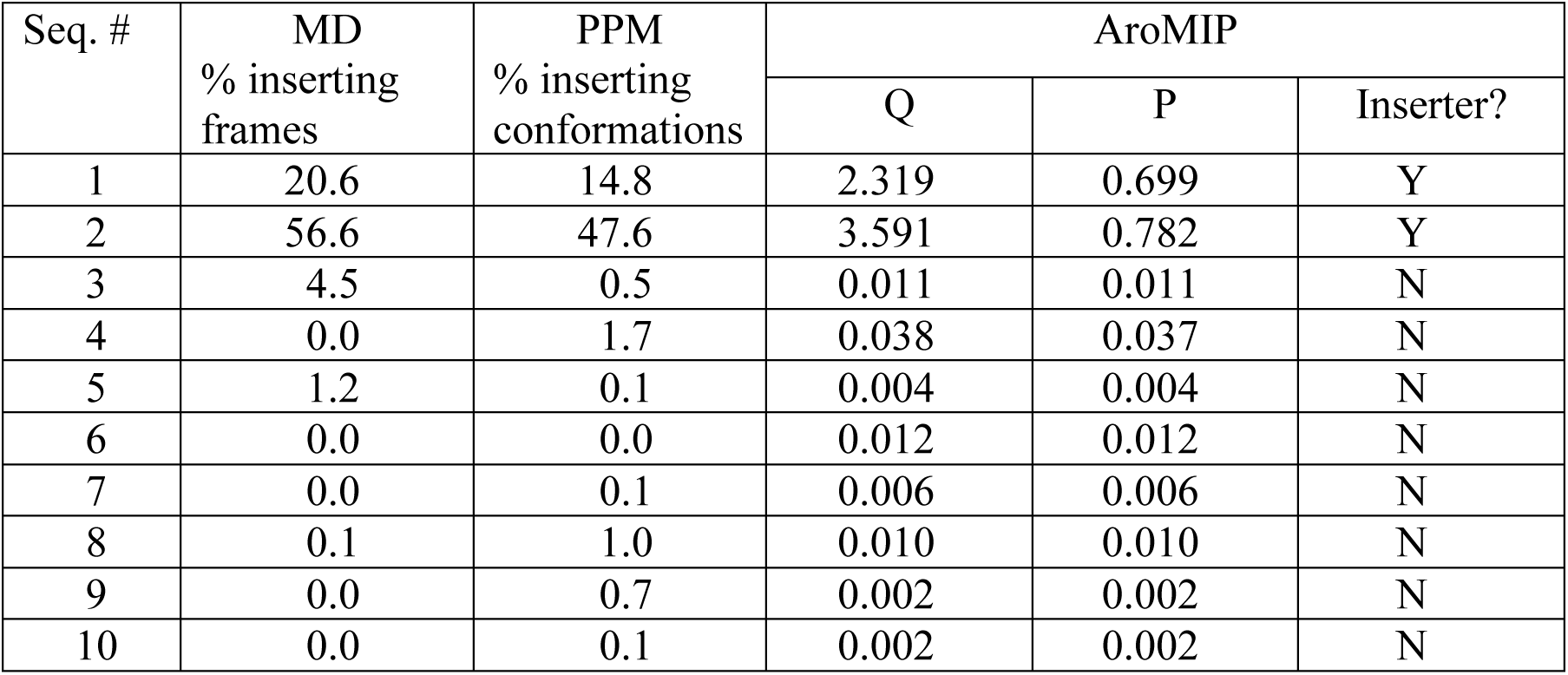
MD, PPM, and AroMIP results on the initial dataset.

The simulations of sequences 1-3 revealed distinct membrane-bound states and pathways to membrane insertion (see Supporting Text 1 for more details). For the F- and W-centered peptides (sequences 1 and 2), a peak of the aromatic Z_tip_ distributions at −6.5 Å defines the inserted state (Fig. 2A, B), where the aromatic sidechains form hydrophobic interactions with acyl chains. The aromatic ring is mostly oriented parallel to the membrane normal, but can also turn sideways (Fig. S1A-C). In both populations, the indole nitrogen atom of W can hydrogen bond with glycerol oxygen atoms. In comparison, the Y-centered sequence 3 features a partially inserted peak at Z_tip_ = −2.5 Å, where the sidechain hydroxyl can hydrogen bond with glycerol oxygen atoms (Fig. 2C) and, correspondingly, the Y sidechain and the glycerol C2 carbon atom of a partner lipid rise and fall together (Fig. S1E, F). For each peptide, a second peak in aromatic Z_tip_ constitutes an intermediate state. Figure 2D illustrates the aromatic sidechains in their respective (partially) inserted states. The aromatic sidechains sample either exclusively the (partially) inserted state (Fig. S2A, B, red curves; Fig. S2C, solid red curve) or both the intermediate and (partially) inserted states (Fig. S2A, B, insets; Fig. S2C, dashed red curve). They have a tendency to form cation-π interactions with a lipid choline group, especially at their respective intermediate states (Fig. 2A-C), occurring in 13.8%, 26.4%, and 24.2% of all the membrane-interacting frames for sequences 1 to 3, respectively (Fig. S2).

**Fig. 2.** Differences and similarities in membrane insertion and cation-π interactions among F-, W-, and Y-centered peptides. (A) Aromatic Z_tip_ distribution for an F-centered peptide (sequence 1). Insets: snapshot with F deeply inserted to interact with lipid acyl chains (bottom), or above the membrane to form cation-π interaction with a choline group (top). (B) Aromatic Z_tip_ distribution for a W-centered peptide (sequence 2). Insets: snapshot with W deeply inserted to interact with lipid acyl chains and form a hydrogen bond with a glycerol oxygen atom (bottom), or in the headgroup region to form cation-π interaction with a choline group (top). (C) Aromatic Z_tip_ distribution for a Y-centered peptide (sequence 3). Insets: snapshot with Y in the headgroup region to form a hydrogen bond with a glycerol oxygen atom (bottom), or near the membrane to form cation-π interaction with a choline group (top). In (A-C), results were accumulated from the last 100 ns of 20 simulations of each peptide; a red dashed line is drawn at Z_tip_ = −3.1 Å. (D) Illustration of stable positions of F, W, and Y with respect to the membrane. (E) Extremely strong correlation between the percentage of membrane-inserting frames in MD simulations and the percentage of membrane-inserting conformations in PPM runs (correlation coefficient = 0.995).

Membrane insertion is not an act of the central aromatic sidechain alone. Rather, it is helped or hindered by the flanking residues. Figure S3A, B displays the Z_tip_ coordinates of all sidechains in F-centered sequence 1 and W-centered sequence 2 in the last 100 ns of the inserted simulations; Fig. S3C displays the corresponding results for Y-centered sequence 3 in the two partly inserting simulations. For all three sequences, the central aromatic sidechain is consistently inserted deeper than flanking sidechains, as shown by the average over replicate simulations (Fig. S3, solid black curves), justifying our focus on aromatic residues. For F-centered sequence 1 (Fig. S3A), the inserted state is stabilized by intermittent penetration of I[+2] (two residues upstream of the central aromatic residue) into the acyl chain region and snorkeling of R[+4], which means that its aliphatic portion contacts acyl chains while its terminal guanidinium group interacts with phosphates. Additional stabilization is provided by salt-bridge formation with lipid phosphates by all the basic sidechains. In the case of W-centered sequence 2 (Fig. S3B), membrane penetration by M[+3] and L[+4] and snorkeling and salt-bridge formation by R[+1] are the major stabilizing factors for the inserted state. For Y-centered sequence 3 (Fig. S3C), membrane penetration by L[+3] and salt-bridge formation by R[-3] reinforce membrane binding.

Inspecting the entire trajectories of sequences 1-3 (illustrated by Fig. 1B), we arrive at the following pathways for membrane insertion of aromatic-centered motifs. The pathways generally consist of three steps: initial membrane contact, further stabilization at the membrane surface, and insertion by the central aromatic sidechain (Fig. 3). These steps lead to poses illustrated and labeled as 1, 2, and 3 in circles in Fig. 3; poses 2 and 3 correspond to the intermediate and (partially) inserted states, respectively, in Fig. 2A-C. Pose 3 is already described in the preceding paragraph; we now describe poses 1 and 2 (Fig. S4). For F-centered sequence 1, R[-4], R[-3], and K[-1] make the initial membrane contact, via long-range electrostatic interactions with phosphates, especially those on acidic lipids (Fig. S4A; Fig. 3A pose 1). R[+4] then tethers the opposite end of the peptide to the membrane, priming the F sidechain for cation-π interaction with a choline (Fig. 3A pose 2). For W-centered sequence 2, R[+1] makes the initial membrane contact, followed by membrane penetration by L[+4] (or M[+3]) (Fig. S4B; Fig. 3B pose 1). With deeper penetration by L[+4] and M[+3], the W sidechain now forms cation-π interaction in the headgroup region (Fig. 3B pose 2). For Y-centered sequence 3, the initial membrane contact can occur through R[-3] via electrostatic interaction or the Y sidechain itself via cation-π interaction (Fig. S4C; Fig. 3C pose 1). Then, R[-3] and L[+3] pin the peptide to the membrane, placing the Y sidechain for cation-π interaction with a choline (Fig. 3C pose 2).

**Fig. 3.** Three-step pathways to membrane insertion. (A) F-centered sequence 1. Basic sidechains at positions −4, −3, and −1 first tether one end of the peptide to the membrane surface. R[+4] then tethers the peptide at the other end, situating the central F for cation-π interaction. Finally, the aromatic sidechain is inserted into the acyl chain region. (B) W-centered sequence 2. The R[+1] sidechain initiates membrane contact. With possible stabilization by the membrane penetration of an aliphatic residue at position +3 or +4, the central W forms cation-π interaction with a choline and then enters the acyl chain region. (C) Y-centered sequences. The initial membrane contact can be formed by either a flanking basic sidechain (R[-3]; via electrostatic interaction) or the central Y itself (via cation-π interaction). After stabilizing at the membrane surface by R[-3] and L[+3], the central Y lowers into the headgroup region to form a hydrogen bond with a glycerol oxygen atom. The aromatic ring occasionally dips into the acyl chain region, bringing the lipid glycerol with it and producing local curvature.

Previous MD and NMR studies using longer constructs provide validation of our all-atom simulations using 9-residue fragments. Our F-centered sequence 1 was taken from the disordered juxtamembrane domain (residues 440-465) of TrkA; Wang et al.^13^ used all-atom MD simulations to characterize the insertion of this 26-residue region into membranes with a simplified representation. The insertion distance profile in that study is quantitatively reproduced by our simulations (Fig. S5), both showing the deepest insertion by the central F residue. Likewise, our W-centered sequence 2 was taken from the disordered variable domain (residues 497-607) of Drp1. Solution NMR data of Mahajan et al.^5^ on this 111-residue region indicated that residues **W**_552_RG are partially helical and insert into cardiolipin-containing membranes. In one simulation of sequence 2, **W**RGML form an α-helix lying parallel to the membrane surface (Fig. S6A), much like an amphipathic helix. In other membrane-inserting simulations, sequence 2 does not form an α-helix (Fig. S6B, C), showing that the helical structure is only partially stable. When averaged between simulations with helical and non-helical structures, the secondary chemical shifts have similar magnitudes (0.2 to 2 ppm; Fig. S6D) to those reported by Mahajan et al.

### PPM provides a fast alternative to all-atom MD simulations in separating inserters and non-inserters

As demonstrated above, all-atom MD simulations are very effective in separating membrane inserters from non-inserters and providing mechanistic insights, but they are time-consuming and therefore not feasible to apply to a very large set of sequences in order to reach a comprehensive understanding on the contributions of flanking residues to membrane insertion. To address this limitation, we developed a fast workflow based on the PPM method,^52^ originally developed for membrane placement of structured proteins. We adapted this method for disordered peptides, by using a batch of random conformations generated by the TraDES algorithm^54^ to represent each peptide (Fig. 4A). Each PPM run, using a peptide conformation as input, produced a pose of the peptide with respect to the membrane (Fig. 4B) and a membrane interaction free energy, ΔG. Figure 4C displays a scatter plot of ΔG and aromatic Z_tip_ data for a batch of 100 conformations of the F-centered sequence 1. We choose a cutoff of −1.5 Å to label a peptide conformation as membrane-inserting. This cutoff is less strict than in the all-atom MD simulations, due to the rigid treatment of peptides and implicit treatment of the membrane by PPM, but we compensate by adding a cutoff on ΔG, at −4.1 kcal/mol (Fig. 4C, region bordered by red dashed lines).

**Fig. 4.** Classification of aromatic-centered motifs into membrane inserters and non-inserters by PPM runs. (A) Disordered conformations of 9-residue motifs generated by TraDES. (B) The pose of an F-centered peptide (sequence 1) from a PPM run. The phosphate plane is represented by arrays of small salmon spheres. (C) A scatter plot of ΔG and Z_tip_ data for a batch of 100 conformations of sequence 1. Dash lines represent cutoffs for defining inserters (ΔG ≤ −4.1 kcal/mol and Z_tip_ < −1.5 Å). 15 conformations satisfy the cutoffs. (D) Extremely strong correlation between the percentage of MD simulations classified as membrane-inserting and the percentage of conformations in PPM runs (correlation coefficient = 0.997).

We measure the membrane-insertion status of each peptide by the percentage of membrane-inserting conformations in a batch of 100 conformations, and denote this quantity as Φ. Replicate PPM runs yield highly reproducible results for each of the 10 sequences. The minimum Φ values of sequences 1 and 2 is 11% and 38%, respectively, while the maximum Φ value of sequences 3 to 10 is 3%. Figure S7 displays the mean Φ values and standard deviations for the 10 sequences, showing a clear gap between inserters (sequences 1 and 2) and non-inserters (sequences 3 to 10). Therefore, PPM can unequivocally separate inserters from non-inserters, just like all-atom MD simulations. Indeed, the mean percentage of inserting conformations in PPM (Table 1) shows a very strong correlation with both the percentage of inserting frames in MD simulations (Fig. 2E) and the percentage of MD simulations labeled as membrane-inserting (Fig. 4D). In both cases, the correlation coefficients are very close to 1. This strong correlation firmly supports our use of PPM for fast classification of aromatic-centered motifs into membrane inserters or non-inserters.

To complete the PPM-based workflow, we had to choose a threshold Φ value, Φ_th_, for separating each peptide sequence into inserters and non-inserters. We took into consideration the following factors. First, PPM did not account for lipid composition.^52^ Cell membranes contain various levels of acidic lipids, which tend to attract basic sidechains for membrane association,^7, 16^ thereby facilitating membrane insertion of adjacent aromatic residues,^9, 10^ as also demonstrated in our all-atom MD simulations (Fig. 3A, B). We accounted for the contributions of flanking basic sidechains by lowering Φ_th_. Second, Φ_th_ should fall in the gap between known inserters and non-inserters (Fig. S7). The end result for Φ_th_ is 40 – 15 × R – 10 × K (or 9 if that value is < 9), where R and K are the number of Arg and Lys residues, respectively, in the eight flanking positions of aromatic-centered motif. As a minimum test, applying this threshold to the PPM results (Table 1), sequences 1 and 2 are correctly classified as inserters and sequences 3-10 are correctly classified as non-inserters.

### Expansion to a library of 1.2 × 10^6^ random sequences reveals a membrane insertion code

We next applied the PPM-based workflow to an exhaustive library of sequences where the central position was filled with F, W, or Y and each of the four flanking positions on either side was filled with a reduced amino-acid alphabet comprising L, R, G, N, and E (Fig. 5A). This alphabet was selected to represent the chemical diversity of the 20 natural amino acids: L (Leu) as an aliphatic residue, R (Arg) as a basic residue, G (Gly) standing for the backbone, N (Asn) as a polar residue, and E (Glu) as an acidic residue. The total size of the library is 3 × 5^8^ = 1.2 × 10^6^ sequences.

**Fig. 5.** Flanking residues as determinants of membrane insertion. (A) Illustration of the library of 3 × 5^8^ = 1.2 × 10^6^ sequences. (B) Percentages of membrane inserters and prediction accuracy of AroMIP. (C) L and R are enriched in all flanking positions of membrane inserters. (D) N, E, and G are enriched in all flanking positions of non-inserters. In (C, D), an orange dashed line is drawn at a frequency of 0.2 expected of random distributions. (E) *q* parameters for F-, W-, and Y-centered sequences.

Applying the PPM-based workflow to this exhaustive library, F-centered sequences are found to have the highest membrane insertion propensities, with 48.6% of all the sequences as inserters, followed by W-centered sequences, with 42.4% inserters (Fig. 5B). In contrast, Y-centered motifs show very low membrane insertion propensities, with only 1.3% inserters. This contrast between F/W-centered sequences and Y-centered sequences is consistent with the outcome of the initial set of 10 sequences.

With the large library, we can now examine the distributions of amino acids in flanking positions. Within either the membrane inserters or non-inserters, we calculated the frequencies at which each flanking position was occupied by the five types of amino acids. If the distributions of amino acids in the two subsets were totally random, the occupation frequency of any amino acid at any position would be 0.2. An occupation frequency higher than 0.2 in the inserter subset means that that amino acid promotes membrane insertion. Figure 5C displays the occupation frequencies in the inserter subset. The frequencies have the order L > R > G > N > E. For each amino acid, the occupation frequency is relatively uniform across all eight positions. For L and R, the frequencies in the inserter subset are > 0.2, indicating that these amino acids promote membrane insertion. An aliphatic sidechain like L is favored in the acyl chain region, thereby stabilizing the inserted state (Fig. 3). While burying a charge in the acyl chain region is highly unfavorable, R can snorkel to form both hydrophobic interactions via its aliphatic portion and salt bridges via its terminal guanidinium group (Fig. 3A, B).

The order in occupation frequency is reversed in the non-inserter subset (Fig. 5D), where E, N, and G have values > 0.2, meaning that they suppress membrane insertion. In contrast to R, an acidic sidechain like E is electrostatically repelled by lipid phosphates; its suppression of membrane insertion is thus expected. The suppressive effects of N and G can be attributed to the fact that it is unfavorable to position polar groups, either on the sidechain or the backbone, in the acyl chain region.

Given the much lower membrane insertion propensities of Y-centered sequences relative to F- and W-centered sequences, we report the distributions of amino acids in Fig. S8 for each aromatic center. The results for F- and W-centered sequences (Fig. S8A-D) are similar to the aggregated results shown in Fig. 5C, D. However, for Y-centered sequences, the inserter subset is highly enriched in the insertion promoter L, with corresponding depletion of insertion suppressors E, N, and G (Fig. S8E). Apparently, Y-centered sequences require substantial help from insertion promoters in flanking positions for them to become membrane inserters. In the non-inserter subset of Y-centered sequences, occupation frequencies have only very small differences from the random value of 0.2 (Fig. S8F), reflecting the fact that the non-inserter subset differs from the full set only by a small percentage (i.e., 1.3%) of sequences that belong to the inserter subset.

The PPM results on the exhaustive library clearly demonstrate two simple rules: (1) flanking residues play key roles in determining whether an aromatic-centered motif is a membrane inserter; (2) aliphatic and basic residues promote membrane insertion, whereas polar and acidic residues suppress it. These rules motivated us to test whether a sequence-based method can predict the membrane insertion status. Based on previous models of sequence-dependent properties of IDPs, including membrane-association propensities and drug-interacting residues,^55–58^ we devised a membrane insertion score by multiplying the contributions, *q_i_*, of all flanking residues:

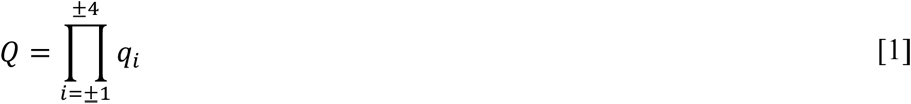

*q_i_* depends on the amino-acid type of the residue at position *i* (the central aromatic residue is at *i*= 0). A; a *q* value > 1 corresponds to promotion of membrane insertion whereas a value < 1 corresponds to suppression of membrane insertion. Note that *Q* has the flavor of a partition function. We then converted this score into a membrane-insertion propensity that ranges from 0 to 1:

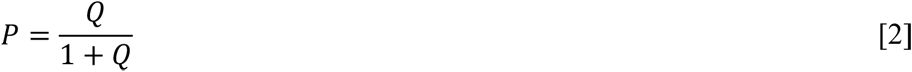

For a given aromatic residue (F, W, or Y), we optimized the five *q* parameters to minimize the difference between *P* and the pre-defined label (0 for non-inserter and 1 for membrane inserter according to PPM) of each sequence. We call this sequence-dependent method AroMIP.

As the training size increases from 1000 to the maximum value 5^8^ = 390625, the *q* parameters quickly converge (Fig. S9). Figure 5E displays the final *q* parameters for F-, W-, and Y-centered sequences. For all three aromatic centers, the *q* values follow the order L > R > G > N > E, as expected from the amino-acid distribution results in the inserter subsets. These values are very similar for F and W centers, > 1 for L and R but < 1 for G, N, and E. The corresponding values for the Y center are lower for all five flanking amino acids. In particular, the *q* value for R dips below 1, making it a weak suppressor of membrane insertion for Y-centered motifs.

As the first test of the sequence-dependent approach, we assessed how well the trained AroMIP model reproduced the PPM-based classification of the 5-amino-acid library, setting a threshold P at 0.5 for inserters (corresponding to *Q* = 1). The retrospective accuracy is 87.5%, 90.1%, and 98.8% for F-, W-, and Y-centered motifs, respectively (Fig. 5B). Such high accuracy levels demonstrate that AroMIP is a promising method.

### AroMIP accurately predicts the membrane-insertion status of IDRs in the human proteome and identifies potential functionally important motifs

We illustrate the utility of AroMIP by applying it to all the IDRs in the human proteome.^53^ From these IDRs, we collected 9-residue motifs centered on an aromatic residue, resulting in a dataset of 24308 motifs (Fig. 6A). We applied the PPM-based workflow to generate labels (membrane inserters or non-inserters) for all these motifs and randomly separated them into a training set (80% of motifs) and a test set (20% of motifs).

**Fig 6.** Implementation of AroMIP on 2.4 × 10^4^ aromatic-centered motifs in IDRs of the human proteome. (A) Illustration of 2.4 × 10^4^ aromatic-centered motifs in IDRs of the human proteome. This dataset was split 80:20 for training and testing, respectively. (B) q parameters for F-, W-, and Y-centered motifs. (C) Performance of AroMIP on the training and test sets.

We then optimized the AroMIP model (equations [1] and [2]) on the training set, now with 20 *q* parameters for the full amino-acid alphabet. Figure 6B displays the resulting *q* parameters for F-, W-, and Y-centered motifs. For F- and W-centered motifs, 12 of the 20 amino acids have *q* > 1 and hence are insertion promoters at flanking positions. This group includes 3 aromatic amino acids (F, W, and Y), 6 aliphatic amino acids (L, I, M, V, P, and A), 2 basic amino acids (R and K), and a single polar amino acid (T). The *q* values of F and W are much higher than those of the other amino acids, indicating unique membrane-insertion propensities. Between F- and W-centered motifs, the *q* values of F and W are higher in the latter; for the remaining 18 amino acids, *q* values are similar between F- and W-centered motifs. In comparison, all 20 *q* values are reduced for Y-centered motifs. Consequently, 5 of the 12 amino acids are no longer in the insertion-promoting group: M, P, T, K, and A. Comparing the *q* values of L, R, G, N, and E here from training on a 20-amino-acid dataset (Fig. 6B inset) against those from a 5-amino-acid dataset (Fig. 5E), we see concordance in several respects, including the same rank order L > R > G > N > E, similarity between F- and W-centered motifs and contrast with Y-centered motifs, and same groupings of insertion promoters and suppressors, except for R in Y-centered motifs, which switches from a weak suppressor in the 5-amino-acid dataset to a weak promoter in the 20-amino-acid dataset. Quantitatively, for these five amino acids, the range of *q* values obtained from the 20-amino-acid dataset is widened somewhat, at both the higher (for L) and lower (for E) ends.

Now let us look at the performance of the trained AroMIP model. In the training set, the percentages of membrane inserters are 59.7%, 51.2%, and 0.7% for F-, W-, and Y-centered motifs, respectively (Fig. 6C). The retrospective accuracy is 91.6%, 91.4%, and 99.4%, respectively. In the test set, the percentages of membrane inserters are 60.1%, 55.0%, and 0.5% for F-, W-, and Y-centered motifs, respectively, which are, as expected, comparable to the counterparts in the training set. The prospective accuracy is 91.2%, 92.0%, and 99.7%, respectively. The latter are essentially identical to the counterparts on the training set, indicating the robustness of the AroMIP model.

We list the top-scoring aromatic-centered motifs in Supplementary Table S1. The F- and W-centered motifs reach the full value of 1 in membrane-insertion propensity, and are typically flanked by two other F and W residues and a combination of aliphatic and basic residues. In contrast, the Y-centered motifs are limited to 0.78 in P, and are flanked by one F or W but a high number of aliphatic residues. Some of these motifs may actually be functionally important. In particular, the transmembrane protein anoctamin-8 resides in the endoplasmic reticulum and tethers the plasma membrane via its C-terminal IDR to assemble Ca^2+^-signaling complexes.^59^ The A_1023_**F**LS**F**K**F**LK_1031_ motif in this IDR may insert into the plasma membrane to facilitate this function. Similarly, GARRE1 localizes to the plasma membrane and intracellular vesicles, likely via a truncated BAR domain, to mediate ciliogenesis.^60^ Membrane insertion by R_885_T**W**P**F**PE**FF**_893_ in the C-terminal IDR may fine-tune this function. Sec16B, with a key role in biogenesis of peroxisomes, tightly associates with endoplasmic reticulum membranes via a central conserved domain but with help from its C-terminal IDR.^61^ The latter may involve membrane insertion by the S_910_G**F**G**WF**S**WF**_918_ motif.

Insertion of an aromatic sidechain into the acyl chain region requires membrane binding of the parent sequence. We asked how well our insertion score (*Q*) is modeled by the membrane binding free energy of the 9-residue motif. As shown in Fig. S10A-C, insertion scores (expressed as −ln *Q*) show moderate to strong correlations with membrane binding free energies (Δ*G*), calculated using the experimentally determined scale for the transfer from water to octanol by Wimley et al.^62^ For F-, W-, and Y-centered motifs in the human proteome, the correlation coefficients range from 0.751 to 0.556. Interestingly, the correlations of our insertion scores with the octanol scale are moderately stronger than with the counterparts with the membrane interfacial scale^24^ (Fig. S10D). This observation can be rationalized because octanol is a mimic of the hydrophobic core instead of the interfacial region,^63^ and hence more in line with our insertion score.

### Additional test cases provide further validation of AroMIP

We have implemented AroMIP as a web server, accessible at https://zhougroup-uic.github.io/AroMIP/, and now report results on additional test cases. First, we go back to our initial set of 10 sequences. The Q and P values for these sequences are listed in Table 1, which result in correct predictions of sequences 1 and 2 as membrane inserters and the other 8 as non-inserters. Moreover, the raw P values are highly correlated with both the percentages of membrane-inserting MD simulations (Fig. S11A) and the percentages of membrane-inserting conformations in PPM runs (Fig. S11B). This correlation suggests that AroMIP can be used not only for binary classification but also for detecting fine differences between motifs.

Next we report the performance of AroMIP on aromatic-centered motifs in 12 other IDPs or IDRs for which information regarding membrane insertion is available (Supplementary Table S2). Using solid-state NMR, Zhang et al.^10^ observed strong cross peaks between acyl chains and five phenol rings from a 25-residue IDR (residues 152-176) of MARCKS. AroMIP predicts very high membrane-insertion propensities for four F residues and high P for the fifth F residue, consistent with the solid-state NMR results. Similarly, solution NMR found cross peaks between the indole ring of W11 in the 86-residue IDP Tat and acyl chains.^44^ AroMIP correctly predicts W11 as a membrane inserter with a high P (= 0.73). In comparison, four other aromatic residues are predicted with lower P values, with one (F38) passing the 0.5 threshold.

Attenuation of NMR signals indicated strong membrane binding by Y511 and weaker binding by Y483 in the C-terminal IDR of Tir.^17^ Whether these Y sidechains were inserted into the acyl chain region was unclear. AroMIP predicts both residues as non-inserters, but with a higher membrane-insertion propensity for Y511 (P = 0.14) than for Y483 (P = 0.01). A slightly higher propensity is predicted for F489 (P = 0.35). Interestingly, these P values correlate with the extents of NMR signal attenuation upon binding DMPG:DHPC membranes. A similar correlation is found for three aromatic residues in the cytoplasmic IDR of mGluR3, with high NMR signal attenuation^16^ and high membrane-insertion propensity (P = 0.72) for F844 and low signal attenuation and low P values (i.e., 0.002 and 0.04) for Y853 and Y861. Moreover, the Y853F mutation increases both the extent of NMR signal attenuation and the membrane-insertion propensity (from 0.002 to 0.33). AroMIP predicts Y853 as a non-inserter, which is consistent with all-atom MD simulations showing that this residue only makes excursions into the acyl chain region.^16^

All-atom MD simulations showed that the aromatic motif **W**_440_**FF**G**W**_444_ in the 57-residue IDR of Bap1 is stably inserted into the acyl chain region.^20^ AroMIP predicts extremely high propensities (close to 1) for these four aromatic residues, due to the presence of flanking F or W (see Fig. 6B). Three other W residues (at positions 420, 432, and 456) are also predicted as inserters (P values range from 0.78 to 0.86); MD simulations did not find membrane insertion by these residues. For KtrB, all-atom simulations showed membrane insertion by F10 in the N-terminal IDR (residues 1-24), but not the adjacent Y11.^9^ In complete agreement, AroMIP predicts a high membrane-insertion propensity (P = 0.75) for F10 but a very low P value (0.07) for Y11. All-atom simulations also found membrane insertion by W21, W25, and W29 in the N-terminal IDR (residues 1-33) of ABHD5, with a greater depth for W25 than for W21 and W29.^49^ AroMIP predicts a perfect P value of 1 for W25, due to the presence of the other two W residues in flanking positions, and somewhat lower values for W21 and W29. Coarse-grained simulations showed membrane insertion by the I_267_**F**PPV_271_ motif in the membrane-proximal IDR (residues 260-280) of the prolactin receptor;^14^ AroMIP predicts a very high membrane-insertion propensity (P = 0.99) for F268.

Cryo-EM structures of OPA1 place W775 and W778 in the membrane-inserting loop deeply into the acyl chain region of cardiolipin-containing membranes.^38, 39^ Likewise, cryo-EM structures suggest that aromatic residues at the tip of the β1–β2 loop in gasdermin D (**W**_48_**FW**_50_) and gasdermin B (**F**_46_**F**_47_) insert into the acyl chain region as an anchor.^40, 42^ This scenario was demonstrated in all-atom MD simulations.^41^ AroMIP predicts very high membrane-insertion propensities for all these aromatic residues. In contrast, a medium propensity (P = 0.55) is predicted for W41 in a loop of PLAT1, consistent with its transient insertion in all-atom MD simulations.^43^

## Discussion

We have used three complementary approaches to decipher the membrane insertion code of aromatic-centered motifs in IDPs and IDRs. All-atom MD simulations revealed the characteristics of membrane-bound states and the pathways leading to membrane insertion. F- and W-centered motifs stably insert into the acyl chain region, but a Y-centered motif does so only transiently. Flanking aliphatic sidechains make major contributions to membrane binding and insertion of the motif by penetrating into the acyl chain region; flanking basic sidechains draw the motif to the membrane surface via long-range electrostatic interactions, keep it at the surface via salt bridges, and stabilize the inserted state via salt bridges and snorkeling. The PPM-based workflow enabled a full exploration of sequence space and the discovery that, for membrane insertion by F- and W-centered motifs, flanking F and W are strong promoters, aliphatic sidechains are medium promoters, and R is a modest promoter. Moreover, Y-centered motifs have very low membrane-insertion propensities. The sequence-based AroMIP method accurately captures these rules and enables fast prediction of membrane-insertion status.

The 20 *q* parameters (Fig. 6B) embody the membrane insertion code. For F/W-centered motifs, flanking F and W have very high *q* values, highlighting their unique ability in membrane insertion and again justifying our focus on aromatic-centered motifs. Consequently, F/W clusters result in very high membrane-insertion propensities. This pattern is seen in the top-scoring F/W-centered motifs from IDRs of the human proteome (Supplementary Table S1), in the **W**_440_**FF**G**W**_444_ motif of Bap1, the **W**_21_LTG**W**LPT**W**_29_ motif of ABHD5, and the **W**_48_**FW**_50_ motif of gasdermin D. For Bap1, mutating WFFG to LGPE significantly reduced membrane binding.^20^ Similarly, mutating W48 and W50 to E (or G in the mouse homolog) rendered gasdermin D defective in pore formation.^40, 64^ It will be interesting to see whether the predicted effects of F/W clusters in the C-terminal IDRs of anoctamin-8, GARRE1, and Sec16B pan out. Whereas F/W clustering can ensure membrane insertion, aliphatic and basic residues can tune the membrane-insertion propensity to desired values. Aliphatic and basic residues can regulate membrane insertion in distinct ways. While aliphatic residues have a somewhat higher ability to stabilize the inserted state, basic residues can draw the aromatic-centered motif to the membrane surface prior to insertion.

The most unexpected result is the enormous difference in membrane-insertion propensity between Y and F or W. In both the exhaustive 5-amino-acid library and the dataset from IDRs of the human proteome, F/W-centered motifs have ∼50% chance of being a membrane inserter, but Y-centered motifs have only ∼1% chance of being an inserter (Figs. 5B and 6C). Burial of the tip hydroxyl of Y by itself in the acyl chain region is highly unfavorable; instead, the hydroxyl hydrogen bonds with a glycerol oxygen atom, and an excursion of Y into the acyl chain region is accompanied by the glycerol group, creating local curvature in the membrane surface (Fig. 3C). Although the W sidechain also has a polar NH group, its position in the five-membered ring allows it to hydrogen bond with a glycerol oxygen atom while the six-membered ring is buried in the acyl chain region (Fig. 2B). Interestingly, the low membrane-insertion propensity of Y fits well with its roles in signal transduction, where a Y-centered motif has to switch binding partners between a membrane and an effector protein.^11, 15–17^ Stable deep insertion would hinder this switch. Indeed, in many cases, the switch from membrane to effector protein requires phosphorylation of the central Y residue, which weakens membrane binding. Deep insertion would also hamper phosphorylation by limiting the kinase’s access to the Y residue.

We have thoroughly validated AroMIP. The method was motivated by distinct distributions of aromatic, aliphatic, basic, polar, and acidic residues in flanking positions of membrane-inserting aromatic-centered motifs, and quantitatively accounts for their disparate contributions to the membrane-insertion propensity of the central aromatic residue. When tested on IDRs in the human proteome, the accuracy of AroMIP exceeds 90% for F/W-centered motifs and approaches 100% for Y-centered motifs. Additional test cases, previously characterized by experimental or MD studies, provide further validation. We also predict possible functionally important motifs in the IDRs of anoctamin-8, GARRE1, and Sec16B. While these predictions are yet to be tested, they illustrate AroMIP’s value in narrowing the search for key residues in functional studies. AroMIP can likewise guide the design of MD simulation studies, in particular, by focusing attention on putative membrane-inserting motifs. With additional applications of AroMIP and further development, we anticipate that the understanding of membrane insertion will reach the same clarity as for the other modes (via amphipathic helices and polybasic motifs) of IDP-membrane association.

An obvious limitation of the present work is that it is based on a single membrane composition (POPC:POPS:PIP_2_ at 70:25:5 ratio). Cell membranes vary substantially in composition. Based on current understanding of protein-membrane interactions, we speculate that a lower PIP₂ level may reduce the contributions of R and K, and the addition of cholesterol could also decrease deep insertion by increasing bilayer order. Cardiolipin is enriched in the inner mitochondrial membrane and many bacterial plasma membranes; with a net charge of −2, it may also favor the binding of R and K via electrostatic attraction. The preponderance of oxygen atoms in the cardiolipin headgroup makes this lipid a likely hydrogen-bond acceptor of W sidechains.^65, 66^ The conical shape of this lipid may promote membrane defects, which facilitate membrane insertion.^47^ These effects may explain why a W552F mutation reduced the binding affinity of a **W**_552_RG-containing IDR for cardiolipin liposomes by 3.6-fold. A future development of AroMIP will be to capture the effects of membrane composition by membrane-specific parameterization.

Another potential limitation is our use of a 9-residue motif. Including four flanking residues on each side of a central aromatic residue balances model parsimony and capturing the most important contributions of neighboring residues. All-atom MD simulations have shown that different segments of an IDP tend to bind membranes independent of each other, as long-range interactions are relatively rare.^18, 20, 22^ Accordingly, our MD simulation of a 9-residue motif showed very similar results to those from simulations of a longer, 26-residue IDR by Wang et al. (Figure S5). Indeed, even neighboring residues appear to bind membranes independently.^18^ This independence explains why the insertion score in equation [1] is calculated as the multiplication of the contributions of all flanking residues. This formulation is equivalent to Hristova and White’s assumption of additive contributions of all residues when calculating the membrane-binding free energy of peptides.^67^ Indeed, as shown in Fig. S10, our insertion scores and their binding free energies show moderate to strong correlations. Nevertheless, the current AroMIP neglects long-range or polyvalent interactions, transient secondary structure formation, and post-translational modifications. These factors will also be targets of future developments.

## Methods

### MD simulations

Each of the ten nine-residue peptides from the initial dataset in a disordered conformation generated through TraDES^54^ was placed near a membrane, with the central aromatic residue just above the phosphate plane. The peptides were N-terminally capped with an acetyl group and C-terminally capped with an amide group to mimic the state of the motifs within the full-length protein. The upper leaflet of the membrane was composed of 70 POPC, 25 POPS, and 5 PIP_2_ lipids; the lower leaflet was composed of 97-99 POPC lipids, with matching surface areas between the two leaflets. The system was then solvated in a rectangular box (with dimensions listed in Supplementary Table S3) with water and NaCl (neutralizing ions plus 150 mM) using the CHARMM-GUI web server.^68^ The force fields were CHARMM36^69^ for proteins and TIP3P^70^ for water. After energy minimization for 10000 steps, each system was equilibrated in six steps in NAMD 3.0.^71^ The first two were at constant temperature and volume, while the remaining four were at constant temperature and pressure. The timesteps were 1 fs in the first three and increased to 2 fs in the last three steps. The simulation times were 125 ps in the first three and 500 ps for the last three steps. During these six steps, restraints on the lipid phosphorus atoms were gradually reduced with the force constant changing from 5 kcal/mol/Å^2^ to 0; similarly, the force constant on peptide heavy atoms was reduced from 10 kcal/mol/Å^2^ to 0. Production runs, with 20 replicates for each peptide, were carried out at constant temperature and pressure without restraints and a 2 fs timestep using pmemd.cuda^72^ in AMBER 22^73^, for a simulation time of 1 µs.

Bond lengths containing hydrogen were constrained using the SHAKE algorithm.^74^ Long-range electrostatic interactions were treated using the particle mesh Ewald method^75^ with a nonbonded cutoff of 12 Å. The temperature was maintained at 300 K by the Langevin thermostat^76^ with a damping constant of 1 ps^-1^; the pressure was maintained at 1 bar by the Berendsen barostat.^77^ Snapshots were saved every 100 ps for analysis.

### MD and PPM data analysis

Z_tip_ distances were calculated for the last 100 ns of simulations using a custom Python code by measuring the difference between the Z coordinate of the tip carbon atom of an aromatic sidechain and the average Z coordinate of phosphorus atoms in the upper leaflet. Frames in which the peptide moved to the image membrane (Z_tip_ beyond the midpoint between the upper leaflet and its periodic image) were excluded from analysis. Distributions were calculated using only the remaining frames. The tilt angle of the aromatic sidechain was the polar angle of a vector connecting two diagonal atoms on the six-membered ring (Cγ-Cζ in F and Y and Cδ_2_-Cη_2_ in W; Fig. S2A). A peptide was considered membrane-interacting when any of its heavy atoms was within 5 Å of any heavy atom of the membrane. Cation-π interaction between a lipid headgroup and an aromatic sidechain was defined when the distances between a choline nitrogen and carbon atoms on the six-membered ring in the aromatic sidechain were less than 7 Å and these distances differed by less than 1.5 Å (https://github.com/weberdak/flexible-interaction-tool/blob/master/flirt.tcl; code written by D. K. Weber).^78^ Cα chemical shifts were calculated every 1 ns over the last 100 ns of simulations using SHIFTX2.^79^ After averaging the 100 frames in a simulation and then over three selected simulations, the corresponding random-coil values were subtracted to yield Cα secondary chemical shifts.

PPM runs used the source code from https://console.cloud.google.com/storage/browser/opm-assets/ppm3_code,^52^ with a batch of 1000 or 100 conformations generated by TraDES as input for each peptide. The output for each peptide conformation was the membrane interaction free energy (ΔG) and a coordinate file representing the peptide’s pose relative to the membrane. Here Z_tip_ was defined using the most buried ring atom on the central aromatic residue.

### Aromatic-centered dataset from IDRs of the human proteome

All IDR sequences in the human proteome were collected from https://mobidb.org/.^53^ Each IDR sequence was then scanned for 9-residue motifs centered at F, W, or Y; duplicates were removed, yielding a dataset of 24308 sequences.

### AroMIP implementation

For each peptide sequence, we calculated a membrane insertion score by multiplying the contributions of all flanking residues (see equation [1]). We then converted the score to a membrane insertion propensity ranging from 0 to 1 (see equation [2]). For a given aromatic residue (F, W, or Y), the *q* parameters were optimized by using the scipy.optimize module^80^ in Python to minimize the difference between the predicted inserted propensity and the pre-defined label (0 for non-inserter and 1 for membrane inserter) for sequences in a training set. The resulting AroMIP model was then applied to a test set for reporting accuracy and implemented as a web server. A threshold propensity of 0.5 separated membrane inserters from non-inserters.

## Acknowledgement

This work was supported by National Institutes of Health Grant GM118091.

## Notes

### Competing Interest Statement

The authors have declared no competing interest.

### Summary of Updates

A new supporting figure added; two paragraphs added in Discussion.

