## Supporting text, tables, and figures for "A membrane insertion code for intrinsically disordered proteins"

*Supplementary Information*

**Supporting Text 1**

We further analyzed the simulations of sequences 1-3 to characterize membrane-bound states and pathways leading to membrane insertion. For both the F- and W-centered peptides, the aromatic $Z_{tip}$ distributions in the last 100 ns of all 20 simulations feature a peak at -6.5 Å (corresponding to the inserted state; Fig. 2A, B), where the aromatic sidechains form hydrophobic interactions with acyl chains. According to the tilt angle of the aromatic ring (Fig. S1A), the inserted state splits into two populations (Fig. S1B, C). The major population has the aromatic ring close to parallel to the membrane normal (tilt angle ~ 150°), surrounded by acyl chains (Fig. S1B, C, snapshots on the right). In the minor population, the aromatic ring turns sideways (tilt angle ~ 90°), with the surrounding acyl chains also bent to maintain hydrophobic contact (Fig. S1B, C, snapshots on the left). In both populations, the indole nitrogen atom of W can hydrogen bond with glycerol oxygen atoms. Another difference between F-centered sequence 1 and W-centered sequence 2 is that the latter has a second peak at $Z_{tip}$ = -2.5 Å, in the headgroup region, whereas the former has a second peak at $Z_{tip}$ = 7.5 Å, above the membrane. In comparison, for the Y-centered sequence 3, the peak with the lowest $Z_{tip}$ is at -2.5 Å (above the cutoff for membrane insertion and termed partially inserted), where the sidechain hydroxyl can hydrogen bond with glycerol oxygen atoms (Fig. 2C). In the subpopulation where the aromatic ring is inserted into the acyl chain (with tilt angle ~ 130°), a neighboring lipid is also lowered, such that the hydroxyl-glycerol hydrogen bond is maintained (Fig. S1D, E). Indeed, the Y sidechain and the glycerol C2 carbon atom of the partner lipid rise and fall together (Fig. S1E), with a strong correlation between the respective $Z_{tip}$ and Z coordinate (correlation coefficient at 0.78; Fig. S1F). The Y-centered sequence 3 also has a second peak, at $Z_{tip}$ = 2.5 Å (Fig. 2C). For each peptide, the second peak in aromatic $Z_{tip}$ constitutes an intermediate state.

Figure 2D compares the poses of aromatic sidechains in sequences 1-3 in their respective (partially) inserted states. In the last 100 ns of the membrane-inserting simulations of sequences 1 and 2, the F and W sidechains sample only the inserted state (Fig. S2A, B, red curves). In other, partly inserting simulations, the F and W sidechains sample both the intermediate and inserted states (Fig. S2A, B, insets). Of the two partly inserting simulations of Y-centered sequence 3, the aromatic sidechain is confined to the partially inserted state in the simulation with more inserting frames (57.0%) but samples both the intermediate and partially inserted states in the other simulation (Fig. S2C, solid and dashed red curves). At their respective intermediate states, all three aromatic sidechains have a tendency to form cation-π interactions with a lipid choline group (Fig. 2A-C). Cation-π interactions are formed in 13.8%, 26.4%, and 24.2% of all the membrane-interacting frames in the 20 MD simulations of sequences 1 to 3, respectively (Fig. S2).

**Supplementary Table S1. Top scoring motifs in IDRs of the human proteome**

| Uniprot ID | Protein name | Sequence | # of aromatic | # of aliphatic[a] | # of basic | Membrane-insertion propensity |
|---|---|---|---|---|---|---|
| F-centered | | | | | | |
| K7EN89 | Midnolin | RFILFKRPW | 3 | 3 | 3 | 1.000 |
| Q70EL4 | UBP43 | LFSRFLLAL | 2 | 5 | 1 | 1.000 |
| H3BN95 | INTS14 | FPLPFPFPS | 3 | 5 | 0 | 1.000 |
| Q70EL4 | UBP43 | LRRLFSRFL | 2 | 3 | 3 | 0.999 |
| Q96JE7 | Sec16B | GFGWFSWFR | 5 | 0 | 1 | 0.999 |
| K7EQI6 | ARHGAP33 | LLPFFPHMP | 2 | 6 | 0 | 0.999 |
| E9PJ49 | CCKBR | LMPVFLIPR | 1 | 7 | 1 | 0.999 |
| H0Y542 | BCAS1 | LGLAFRKFF | 3 | 3 | 2 | 0.999 |
| Q9HCE9 | Anoctamin-8 | AFLSFKFLK | 3 | 3 | 2 | 0.999 |
| O15063 | GARRE1 | RTWPFPEFF | 4 | 2 | 1 | 0.999 |
| W-centered | | | | | | |
| Q9UHK0 | NUFIP1 | SWMFWAMLP | 3 | 5 | 0 | 1.000 |
| Q96JE7 | Sec16B | SGFGWFSWF | 5 | 0 | 0 | 1.000 |
| H3BUN7 | SH2B1 | PWLSWSPWL | 3 | 4 | 0 | 1.000 |
| O75807 | GADD34 | FLKAWVYWP | 4 | 4 | 1 | 1.000 |
| H0YFY4 | HMGA2 | RPRKWWLLM | 2 | 4 | 3 | 0.999 |
| KAI2556058[b] | COL13A1 | SWASWFTWT | 4 | 1 | 0 | 0.999 |
| Q9BYE0 | hHes7 | PPAFWRPWP | 3 | 5 | 1 | 0.999 |
| H7C1I9 | MAST4 | LLEPWFLPP | 2 | 6 | 0 | 0.999 |
| Q9Y4K4 | MEKKK 5 | FMLQWNPFV | 3 | 4 | 0 | 0.999 |
| H0YFY4 | HMGA2 | PRKWWLLMK | 2 | 4 | 3 | 0.999 |
| Y-centered | | | | | | |
| A0PJX2 | TLDC2 | LRWRYTRLP | 2 | 3 | 3 | 0.778 |
| A0A7P0T883 | C2CD3 | SLLLYPLAF | 2 | 6 | 0 | 0.765 |
| P48634 | PRRC2A | PPFMYPPYL | 3 | 6 | 0 | 0.762 |
| P48634 | PRRC2A | MYPPYLPFP | 3 | 6 | 0 | 0.762 |
| A0A1B0GV45 | Myosin XVIIIA | LMMRYLYRP | 2 | 5 | 2 | 0.760 |

[a]Aliphatic residues are: L, I, M, V, P, and A.

[b]GenBank ID.

**Supplementary Table S2. AroMIP predictions on 12 IDPs or IDRs**

| IDP name | Previous study[a] | Sequence[b] | AroMemIn[b, c] |
|---|---|---|---|
| MARCKS<br>(UniProt P29966; residues 152-176) | Solid-state NMR[10] | KKKKKR**F**S**F**KKS**F**KLSG<br>**F**S**F**KKNKK | **F158 0.953**<br>**F160 0.992**<br>**F164 0.956**<br>**F169 0.956**<br>**F171 0.742** |
| Tat<br>(Uniprot P04610; residues 1-86) | Solution NMR[44] | MEPVDPRLEP**W**KHPGSQ<br>PKTACTTCYCKKCCFHC<br>QVC**F**TTKALGISYGRKK<br>RRQRRRPPQGSQTHQVS<br>LSKQPTSQPRGDPTGPK<br>E | **W11 0.727**<br>Y26 0.000<br>F32 0.288<br>**F38 0.509**<br>Y47 0.050 |
| Translocated intimin receptor<br>(Uniprot B7UM99; residues 388-550) | Solution NMR[17] | RRNQPAEQTTTTTTHTV<br>VQQQTGGNTPAQGGTDA<br>TRAEDASLNRRDSQGSV<br>ASTHWSDSSSEVVNPYA<br>EVGGARNSLSAHQPEEH<br>IYDEVAADPGYSVIQNF<br>SGSGPVTGRLIGTPGQG<br>IQSTYALLANSGGLRLG<br>MGGLTSGGESAVSSVNA<br>APTPGPVRFV | W443 0.179<br>Y454 0.024<br>Y474 0.002<br>Y483 0.014<br>F489 0.347<br>Y511 0.140 |
| mGluR3<br>(Uniprot Q14832; residues 839-879) | Solution NMR; all-atom MD simulations[16] | LHLNR**F**SVSGTGTTYSQ<br>SSASTYVPTVCNGREVL<br>DSTTSSL | **F844 0.716**<br>Y853 0.002<br>Y861 0.036 |
| Bap1<br>(Uniprot A0A7Z7YFH0; residues 415-471) | All-atom MD simulations; tryptophan fluorescence[20] | YLGLE**W**KTKTVPYLGVE<br>**W**RTKTVSY**WFF**G**W**HTKQ<br>VAYLAPV**W**KEKTIPYAV<br>PVTLSK | **W420 0.838**<br>Y427 0.061<br>**W432 0.787**<br>Y439 0.289<br>**W440 1.000**<br>**F441 0.998**<br>**F442 0.997**<br>**W444 0.998**<br>Y451 0.139<br>**W456 0.863**<br>Y463 0.263 |
| KtrB<br>(Uniprot O87953; residues 1-24) | All-atom MD simulations[9] | MTQFHQRGV**F**YVPDGKR<br>DKAKGGE | **F10 0.746**<br>Y11 0.065 |

| | | | |
|---|---|---|---|
| ABHD5 (Uniprot Q9DBL9; residues 1-33) | All-atom Gaussian accelerated MD simulations; hydrogen-deuterium exchange[49] | MKAMAAEEEVDSADAGG GSG**W**LTG**W**LPT**W**CPTS | **W21 0.858**<br>**W25 1.000**<br>**W29 0.989** |
| Prolactin Receptor (Uniprot P16471; residues 260-280) | Coarse-grained MD simulations; solution NMR[14] | GYSMVTCI**F**PPVPGPKI KGFD | **F268 0.985** |
| OPA1 (Uniprot O60313; residues 768-782) | Cryo-EM[38, 39] | GPDWKKR**W**LY**W**KNRTQ | **W775 0.999**<br>Y777 0.398<br>**W778 0.990** |
| Gasdermin D (Uniprot P57764; residues 41-53) | Cryo-EM;[40] all-atom MD simulations[41] | VRKPSSS**WFW**KPRYK | **W48 0.996**<br>**F49 0.989**<br>**W50 0.998** |
| Gasdermin B (Uniprot Q8TAX9-1; residues 41-51) | Cryo-EM[42] | GEKRT**FF**GCRH | **F46 0.854**<br>**F47 0.920** |
| PLAT1 (Uniprot O65660; residues 37-49 | All-atom MD simulations[43] | TGSI**W**KAGTDSII | **W41 0.551** |

[a]Numbers in superscript are reference numbers from the main text.

[b]Inserters are indicated by bold letters whereas non-inserters are indicated by an underline.

[c]The value after the residue number is the membrane-insertion propensity.

**Supplementary Table S3. Details of MD simulations in membranes**

| Seq. # | Box dimensions (Å$^3$) | # of atoms |
|---|---|---|
| 1 | 82 x 82 x 103 | 64818 |
| 2 | 81 x 81 x 103 | 64647 |
| 3 | 81 x 81 x 106 | 66106 |
| 4 | 82 x 82 x 101 | 63196 |
| 5 | 82 x 82 x 104 | 64865 |
| 6 | 82 x 82 x 104 | 65384 |
| 7 | 82 x 82 x 103 | 65315 |
| 8 | 81 x 81 x 105 | 64809 |
| 9 | 82 x 82 x 103 | 64150 |
| 10 | 82 x 82 x 104 | 65265 |

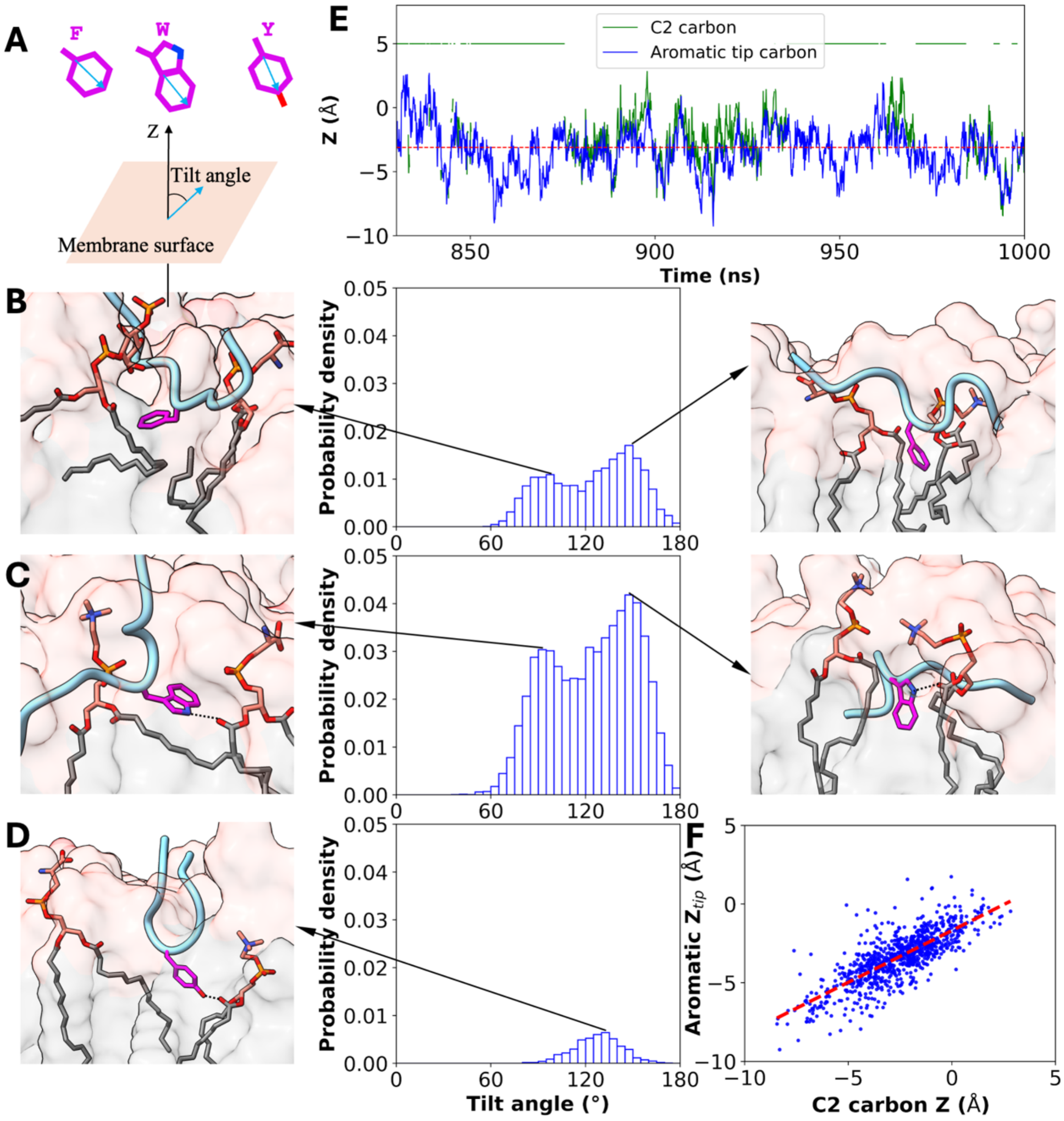


**Fig. S1. Aromatic tilt angle distributions in membrane-inserted frames of MD simulations.** (A) Definition of tilt angle. (B-D) Tilt angle distributions for F-, W-, and Y-centered sequences 1, 2, and 3, respectively, along with snapshots illustrating aromatic poses at major and minor peak angles. (E) Time traces of the aromatic $Z_{tip}$ and Z coordinate of the glycerol C2 carbon atom in a partner lipid in the simulation of Y-centered sequence 3 with 57.0% inserting frames. Lines at Z = 5 Å indicate hydrogen bond breakup; hydrogen bond reformation may be with a different lipid molecule. A red line is drawn at Z = -3.1 Å. (F) Correlation between the aromatic $Z_{tip}$ and Z coordinate of the glycerol C2 carbon atom (correlation coefficient = 0.781).

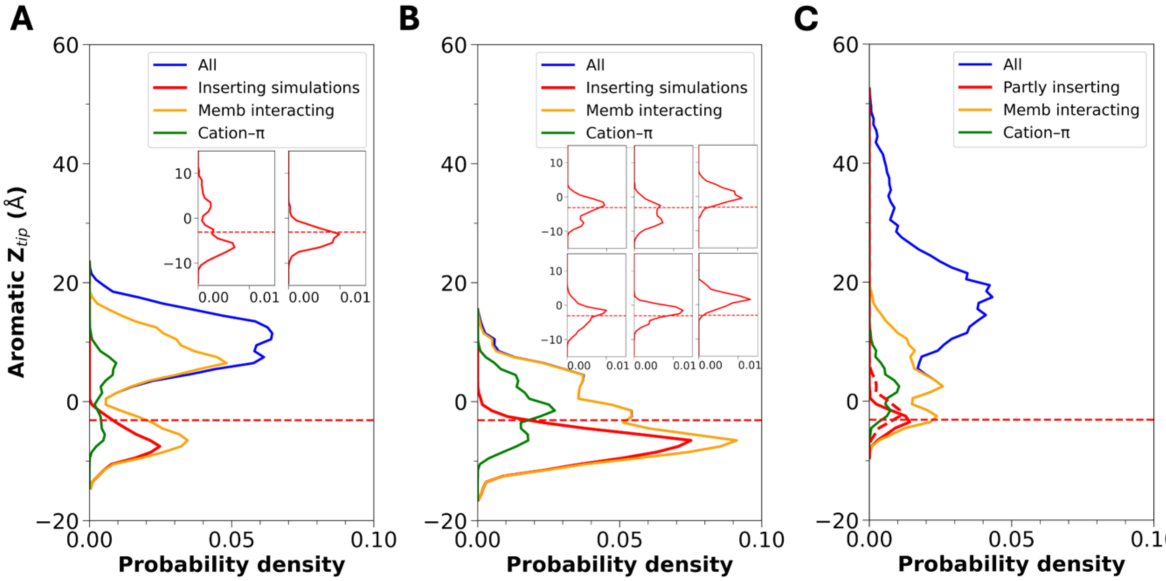


**Fig. S2. Sampling of intermediate and (partially) inserted states and formation of cation-π interactions by aromatic-centered peptides.** (A, B) Aromatic $Z_{tip}$ distributions for F- and W-centered sequences 1 and 2 in all simulations, in membrane-inserting simulations, and among membrane-interacting or cation-π forming frames. Insets: aromatic $Z_{tip}$ distributions in partly inserting simulations. (C) Corresponding results for Y-centered sequence 3, except that partly inserting simulations are presented in the main panel. In all panels, a red line is drawn at $Z_{tip}$ = -3.1 Å.

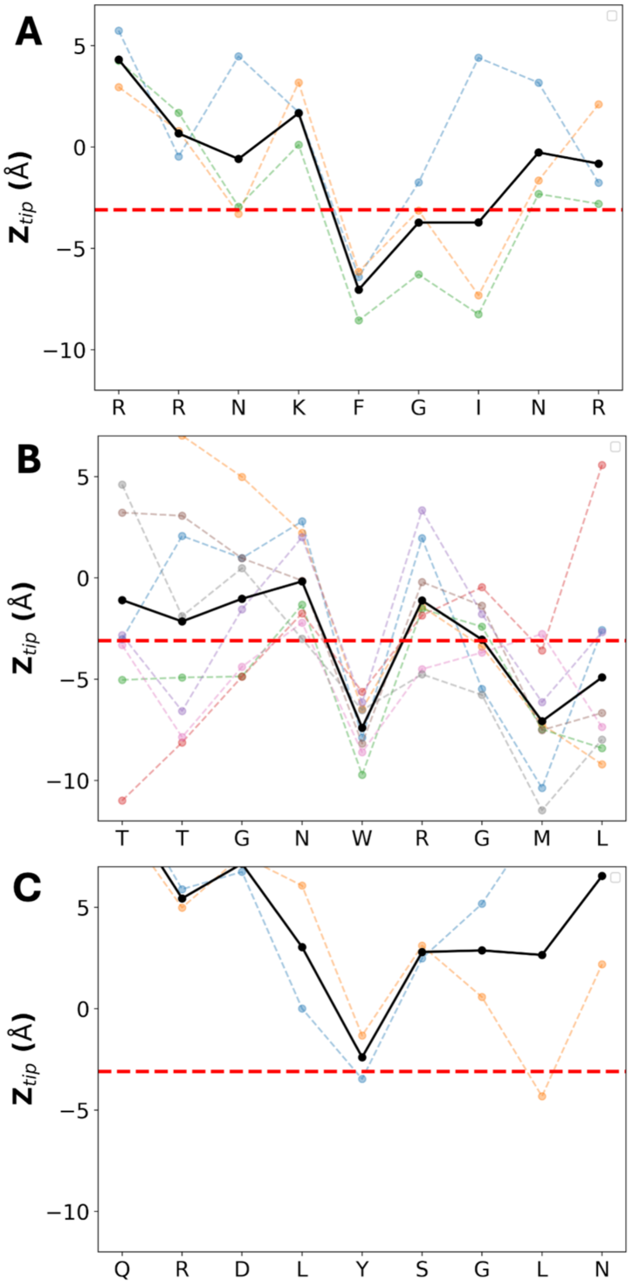


**Fig. S3. $Z_{tip}$ profiles of aromatic-centered peptides calculated from the last 100 ns of (partly) membrane-inserting simulations.** (A, B) Results from 3 and 8 membrane-inserting simulations, respectively, of F- and W-centered sequences 1 and 2. Results from individual simulations are in color; averages among the simulations are in black. (C) Corresponding results for Y-centered sequence 3, except the two simulations are partly inserting. In all panels, a red line is drawn at $Z_{tip}$ = -3.1 Å.

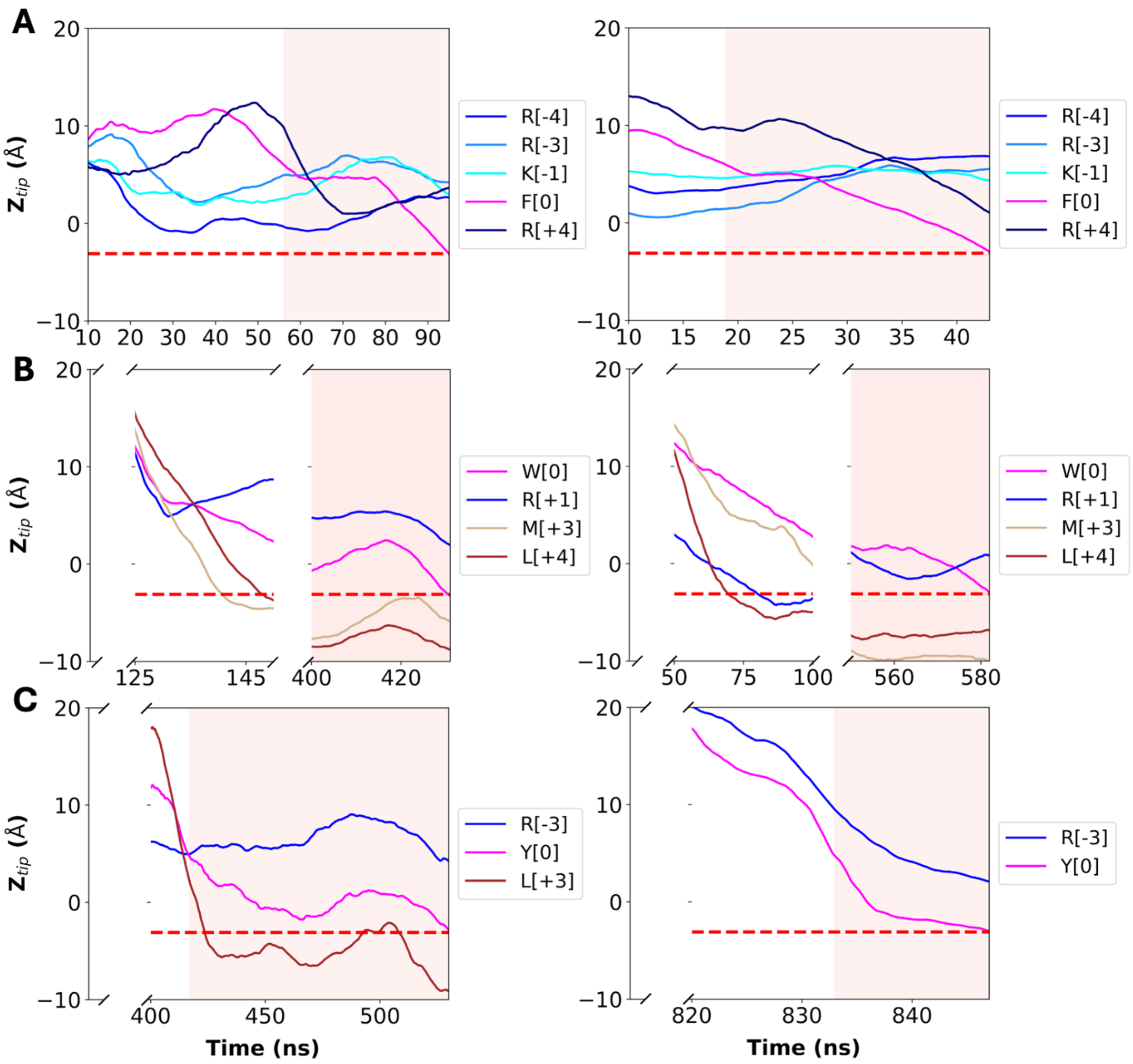


**Fig. S4. Times traces of Ztip coordinates depicting pathways to membrane insertion.** (A-C) Results for F-, W- and Y-centered sequences in two simulations. In (B, C), the peptides initially moved away from membranes and then came back; these initial portions of the simulations are skipped. In addition, in (B) the peptide stayed in the intermediate state for over 200 ns; the initial period in the intermediate state is skipped. Time traces are smoothed using the moving average in a 20.1-ns window. Residues are labeled as X[i], where X is the one-letter code for an amino acid and i is the position relative to the central aromatic residue. In all panels, a red line is drawn at $Z_{tip}$ = -3.1 Å.

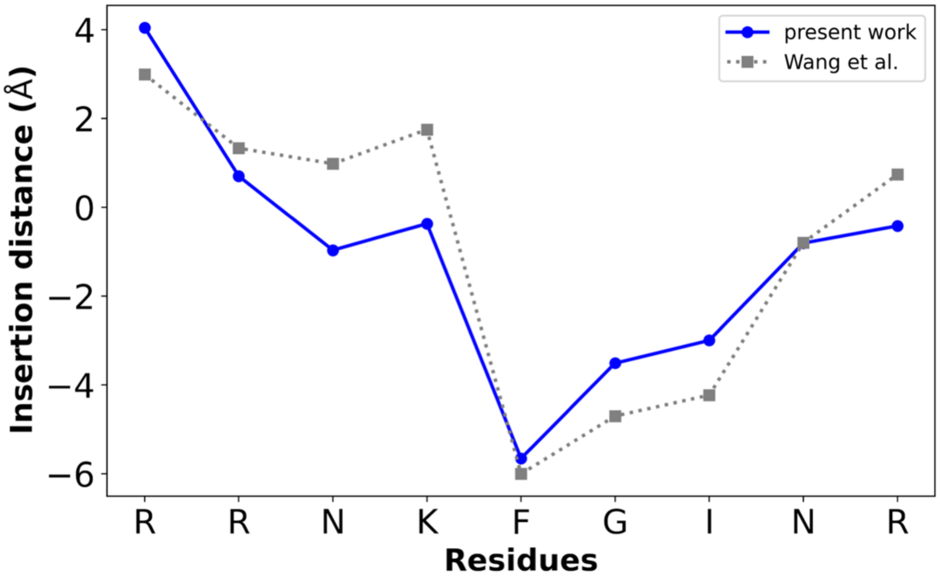


**Fig. S5. Comparison in membrane insertion profile between our simulations with a 9-residue peptide and Wang et al.'s simulations with a 26-residue IDR from TrkA.** The insertion distance was calculated following the protocol of Wang et al., as the difference between the Z coordinate of the all-atom center of mass of a residue and the average Z coordinate of the lipid phosphorus atoms in each frame; the values in the last 50 ns of the three membrane-inserting simulations were pooled and the median is plotted.

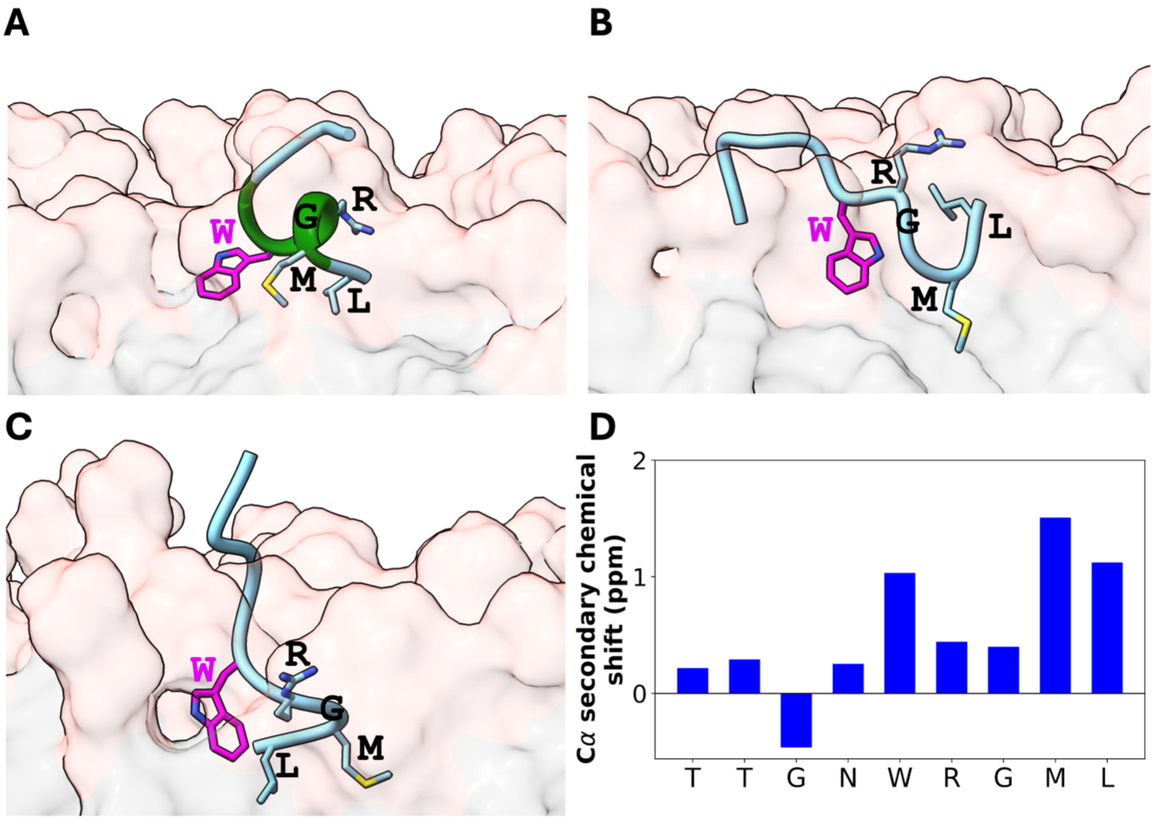


**Fig. S6. Partial helix formation by the WRGML motif of Drp1 in the membrane-inserting simulations.** (A) Snapshot of W-centered sequence 2 in a membrane-inserting simulation, with WRGML forming an amphipathic helix. (B, C) Snapshots in two other membrane-inserting simulations with no helix formation. (D) Cα secondary chemical shifts, averaged over the three membrane-inserting simulations of (A-C).

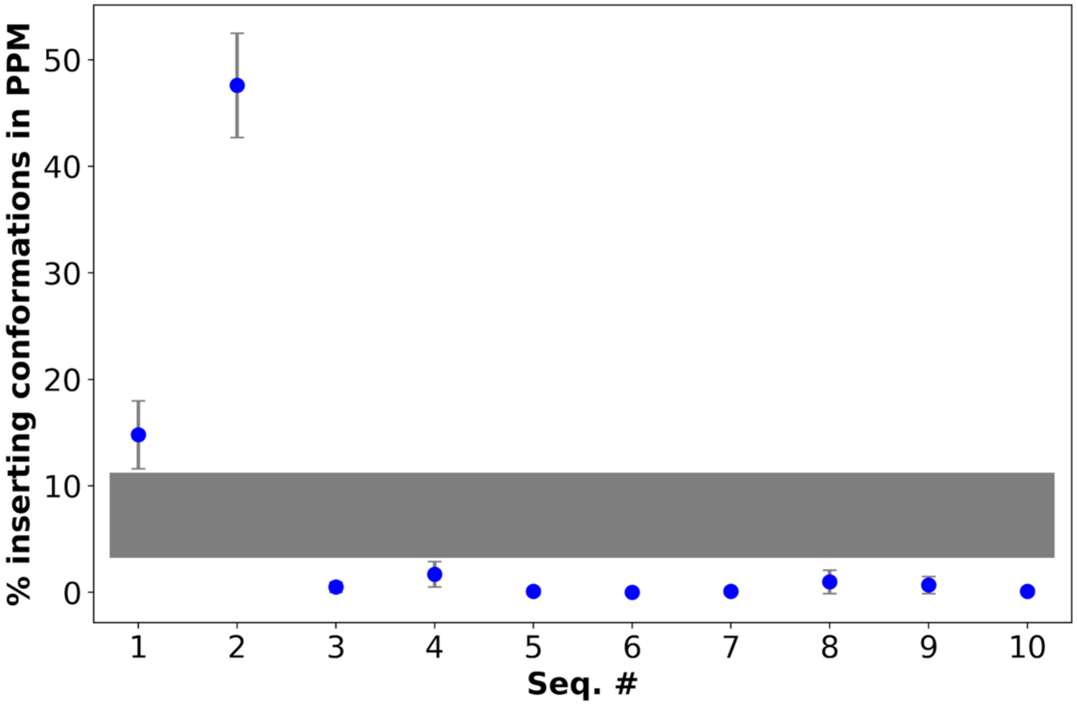


**Fig. S7. Clear separation of membrane inserters and non-inserters by PPM**. For each of the 10 sequences in our initial dataset, the number of conformations in a batch of 100 that satisfy the cutoffs for both aromatic $Z_{tip}$ and $\Delta G$, when averaged over 10 batches, is displayed as a blue circle. Error bars represent standard deviations across the 10 batches. The gray block illustrates the separation between inserters and non-inserters.

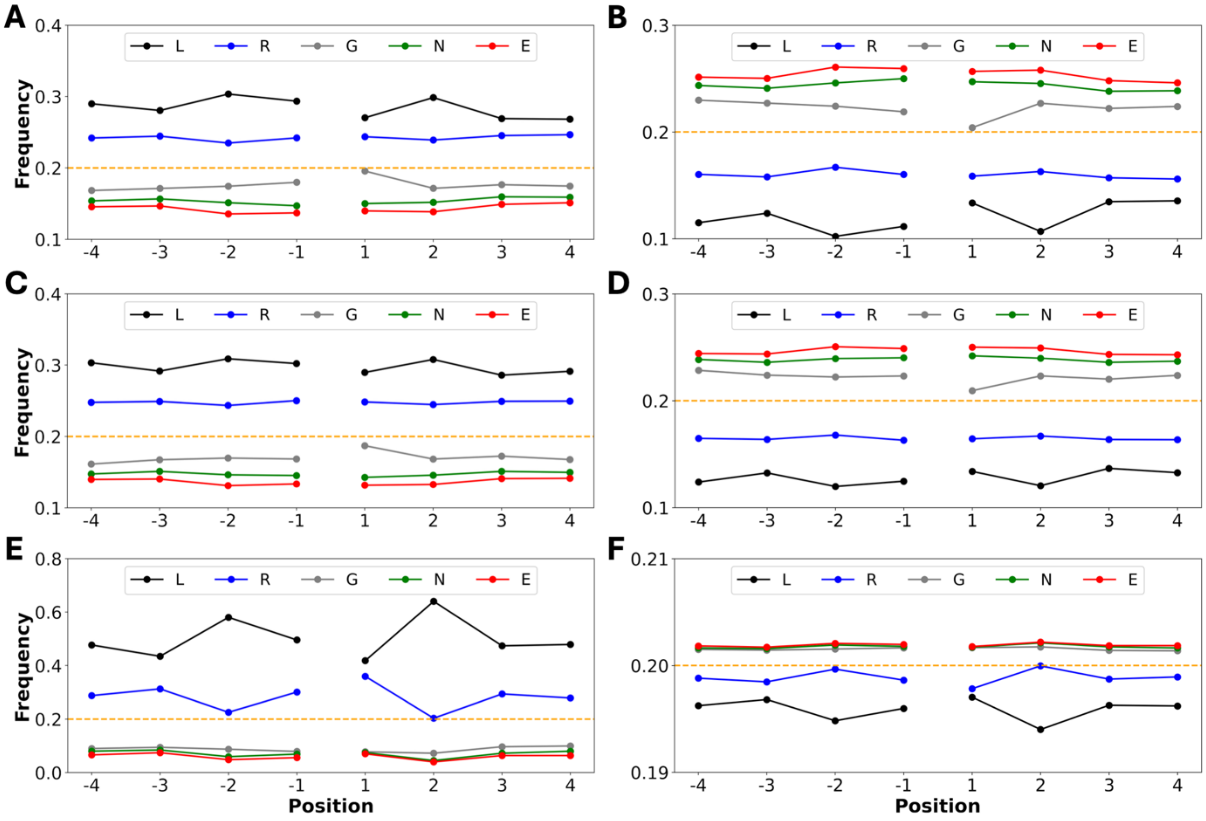


**Fig. S8. Distributions of amino acids in flanking positions of aromatic-centered membrane inserters and non-inserters.** (A, B) F-centered inserters and non-inserters. (C, D) W-centered inserters and non-inserters. (E, F) Y-centered inserters and non-inserters. An orange dashed line is drawn at a frequency of 0.2 expected of random distributions.

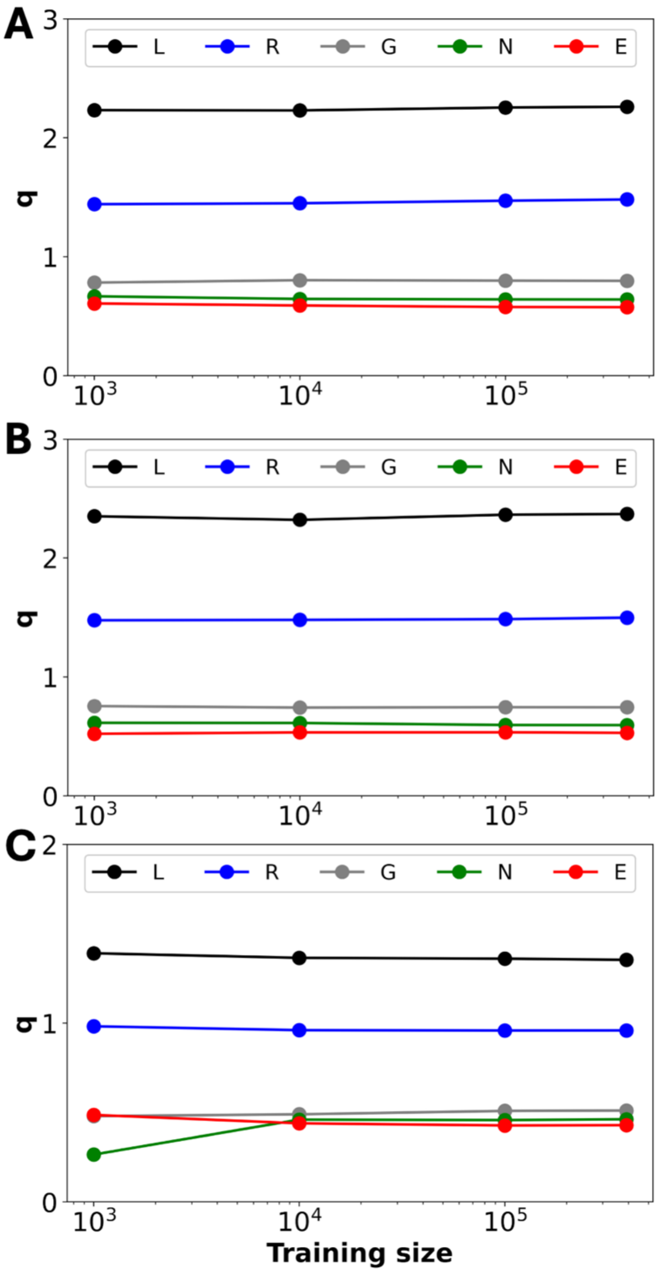


**Fig. S9. Convergence of q parameters at increasing training data size.** (A-C) F-, W-, and Y-centered sequences.

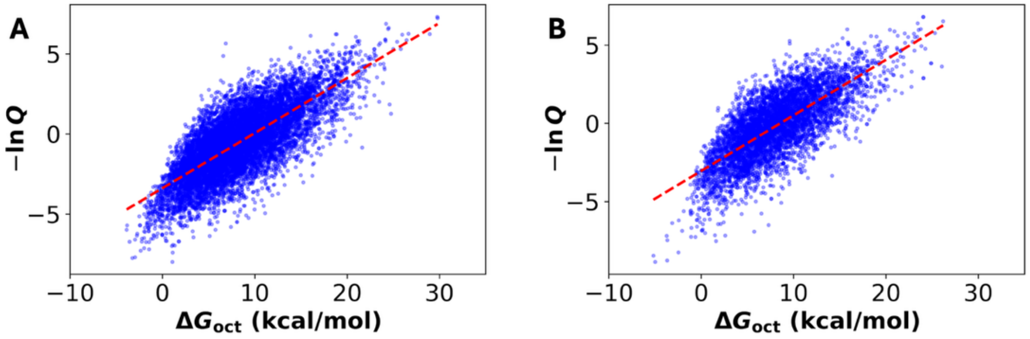


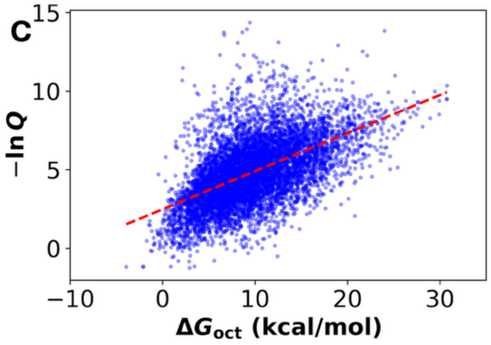


D

| | $\Delta G_{oct}$ | $\Delta G_{if}$ |
|---|---|---|
| **F** | 0.751 | 0.666 |
| **W** | 0.739 | 0.646 |
| **Y** | 0.556 | 0.394 |

**Fig. S10. Correlations of AroMIP insertion scores and membrane-binding free energies for aromatic-centered motifs in the human proteome.** (A-C) Correlations of insertion scores and binding free energies calculated using the octanol scale, for F-, W-, and Y-centered motifs, respectively. (D) Correlation coefficients for two scales: the octanol scale and the membrane interfacial scale.

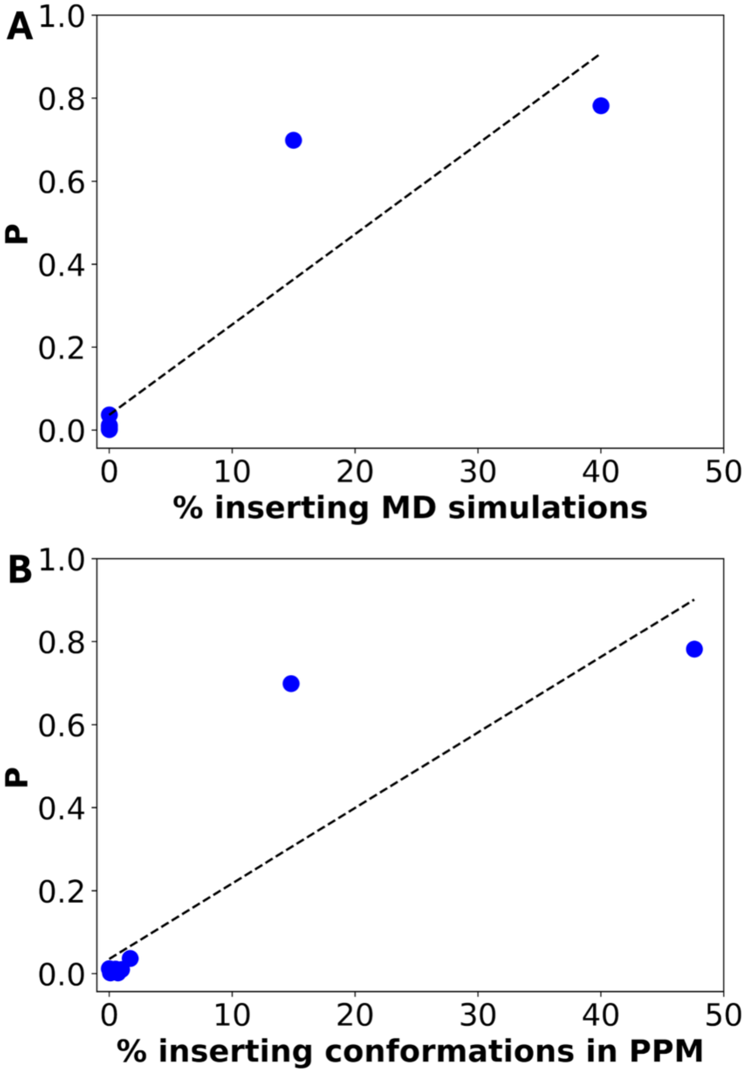


**Fig. S11. Correlations of AroMIP-predicted membrane-insertion propensities and MD and PPM results.** (A) Correlation of membrane-insertion propensity and percentage of membrane-inserting MD simulations (correlation coefficient = 0.918). (B) Corresponding correlation percentage of membrane-inserting conformations in PPM (correlation coefficient = 0.889).
